# Multifaceted influence of pre-mitotic cytotoxicity of primed CD8 T cells on immunity and infection

**DOI:** 10.1101/456392

**Authors:** Durga Bhavani Dandamudi, David A Blair, Raquel Duque do Nascimento Arifa, Juan J Lafaille, Michael L Dustin, Viveka Mayya

## Abstract

Granzyme B mRNA is expressed in primed CD8 T cells within 12 hours, but the consequences of this for the immune response are unknown. We observed that substantial portion of the naïve CD8 T cell repertoire expressed granzyme B and became pre-mitotic cytotoxic cells (PMCs) immediately in response to *Listeria monocytogenes* or *Lymphocytic choriomeningitis virus* infections. The surprising breadth arose from sufficiency of low potency peptide-MHC to induce granzyme B expression in the context of infection. PMCs killed antigen bearing dendritic cells (DCs) in a granzyme B-dependent but largely perforin-independent fashion between 1-2 days post infection. This terminated antigen presentation at 3 days and resulted in reduced clonal expansion. As additional consequences, we highlight that PMCs reduced the burden of DC-borne infectious agents, but also opened a window of vulnerability for secondary infection. Thus, PMCs serve antigen-specific, regulatory and host defence functions, that are innate-like in scale, at the onset of the adaptive immune response.

## Introduction

CD8 T cells form an important arm of adaptive immunity as they are critical for defence against intracellular pathogens and cancerous cells (Masopust et al., 2007; Zhang and Bevan, 2011). Naïve CD8 T cells patrol the secondary lymphoid organs in search of cognate peptide antigen. They recognize antigen presented by dendritic cells (DCs) that have either emigrated from the affected peripheral tissue or are resident in lymphoid tissues and receive soluble or particulate antigens. Following a period of about 24 hours of priming by antigen, costimulatory and inflammatory signals, the naïve CD8 T cells begin to proliferate and differentiate (Kaech and Ahmed, 2001; Mescher et al., 2006). Proliferation leads to clonal expansion and differentiation leads to expression of several effector molecules, adhesion molecules and chemokine receptors over a period of about a week (Best et al., 2013). During the expansion phase, effector CD8 T cells continue to differentiate and can be defined by markers KLRG1 and CD127 as double negative plastic <u>e</u>arly <u>e</u>ffector <u>c</u>ells (EECs) (Obar and Lefrancois, 2010; Plumlee et al., 2015), KLRG1^+^, CD127^-^ terminally differentiated <u>s</u>hort-<u>l</u>ived <u>e</u>ffector <u>c</u>ells (called SLECs) or KLRG1^-^, CD127^+^ long-lived <u>m</u>emory <u>p</u>recursor <u>e</u>ffector <u>c</u>ells (called MPECs) (Joshi and Kaech, 2008). Upon differentiation, the effector CD8 T cells can leave the secondary lymphoid organs and infiltrate peripheral tissues (Zhang and Bevan, 2011). In the affected tissue, they produce chemokines and cytokines to recruit and activate innate immune cells. More importantly, effector CD8 T cells directly kill infected or cancerous cells bearing cognate antigen by triggering their Fas receptor and/or through the release of granzyme B and perforin by exocytosis from specialized granules (Harty et al., 2000).

In contrast to the long-held view that effector function is acquired after clonal expansion, emerging studies demonstrate that effector and regulatory function is initiated even before the first division of activated CD8 T cells. We and others have noted that primed CD8 T cells express multiple cytokines and chemokines within 24 hours of infection or antigen encounter, which precedes the first division (Best et al., 2013; Chiu et al., 2007). Robust but transient expression of inflammatory chemokines by the primed CD8 T cells in this ‘pre-mitotic’ phase was shown to recruit multiple types of immune cells into the reactive secondary lymphoid organ (Sung et al., 2013). Interferon γ (IFNγ) is also produced robustly but transiently by the primed CD8 T cells in this pre-mitotic phase (Curtsinger et al., 2012; Hosking et al., 2014). Blocking IFNγ at this juncture, but not later, severely undermines ensuing clonal expansion of CD8 T cells after <u>l</u>ymphocytic <u>c</u>horio<u>m</u>eningitis <u>v</u>irus (LCMV) infection (Utermohlen et al., 1996) or restricts differentiation into memory cells after *Listeria monocytogenes* infection (Krummel et al., 2018). It is worth noting that all these studies rely on transgenic models that require adoptive transfer of large number antigen specific T cells (~10^6^) to provide sufficient cells for analysis.

Cytolytic molecules are also expressed 12-24 hours after infection (Best et al., 2013; Chiu et al., 2007). Cytotoxicity before the first division has also been documented (Chiu et al., 2007), but was not otherwise distinguished from conventional CTL function. We do not know how cytotoxicity develops in recently primed naïve CD8 T cells and how they kill target cells. Most importantly, the functional relevance of pre-mitotic cytotoxicity in primed CD8 T cells is not known to date. We reveal here that a subset of primed CD8 T cells became granzyme B dependent pre-mitotic cytotoxic cells (PMCs), which are distinct from the post-mitotic conventional cytotoxic T cells (CTLs). We further show that PMCs killed DCs. Elimination of DCs had multiple consequences: 1) reduced burden of DC-borne infectious agents; 2) rapid reduction and termination of ongoing antigen presentation that prevented recruitment of additional antigen-specific naïve T cells; 3) reduced rate of clonal expansion of already primed CD8 T cells; 4) curtailed T cell response to secondary infection. Thus, pre-mitotic cytotoxicity had multiple ramifications to the immune response and course of infection, underscoring the extent to which adaptive immune cells can exert its influence without any clonal expansion, but taking advantage of the degeneracy of TCR recognition to recruit sufficient numbers of cells to have an innate-like scale of response. As highly activated cells become PMCs during priming, they also constitute an auto-inhibitory loop to limit the CD8 T cell response right at the onset. While the conventional effector CTLs and recently primed memory CD8 T cells are also known to kill antigen-presenting DCs in a regulatory feedback (Belz et al., 2007; Yang et al., 2006), PMCs act in a distinct manner very early in parallel with innate defences to protect the host from pathogens and potential immunopathology.

## Results

### Development of pre-mitotic cytotoxicity in primed CD8 T cells

We had previously catalogued genome-wide changes in expression of mRNA during the priming phase in adoptively transferred OTI TCR transgenic CD8 T cells following infection of mice with *L. monocytogenes* expressing chicken ovalbumin (*Lm-ova*) (Best et al., 2013). Granzyme B mRNA was dramatically increased after 12, 24 and 48 hours of priming (Figure 1A), whereas there was no change in perforin (Figure 1A), FasL or Granzyme A (Figure S1A) mRNA levels at these time points. In order to have synchronous activation and clearly defined duration of priming, we had transferred 10^6^ OTI T-cells (OTIs) 24 h after intravenous infection with 5000 *Lm-ova.* We confirmed that T cell activation was indeed more synchronous with this ‘post-infection transfer’ model (Figure S1B), based on CD69 expression (Figure S1C) and cell division (Figure S1D). We established that the first division took place between 28-36 h when OTIs were transferred after infection (Figure 1B). In order to ascertain if *Lm-ova* primed OTIs display pre-mitotic cytotoxicity, we utilized an in vivo cytotoxic assay (Hermans et al., 2004). Ova and control (gp33) peptide-loaded splenocytes, 4 x 10^6^ each as antigen-specific and non-specific target cells, respectively, were co-transferred with 10^6^ naïve OTIs (effector:target, E:T ratio of 1:4) at 24 hours post-infection. Significant reduction in the ova loaded target cells relative to the gp33 loaded target cells was observed 20 h later (Figure 1C). We did not observe any antigen-specific killing activity with 12 hours of priming, but more than 40% of specific targets were killed after 20 hours (Figure 1D). Specific killing activity increased to 63% after 40 hours of priming. This implies that majority of killing happened during the interval between 12-20 hours, when granzyme B was likely at the highest level of expression (Figure 1A). Thus, most of the antigen-specific killing occurs many hours prior to the commencement of the first division. Similarly, pre-mitotic cytotoxicity was also observed with the conventional ‘pre-infection’ transfer of OTIs (Figure S1E). LCMV infection also induced granzyme B expression prior to the first cell division in P14 TCR transgenic T cells that lead to pre-mitotic cytotoxicity (Figure S1F-H). These results, along with previous observations in mice infected with Vaccinia Virus (Chiu et al., 2007) and *Plasmodium yoelii* (Sano et al., 2001), suggest that pre-mitotic cytotoxicity is a wide-spread phenomenon during CD8 T cell priming upon infection. Thus, recently primed CD8 T cells that express granzyme B are referred to subsequently as “PMCs” to distinguish them from conventional cytotoxic T cells, which we will refer to as “CTLs”.

**Figure 1:**
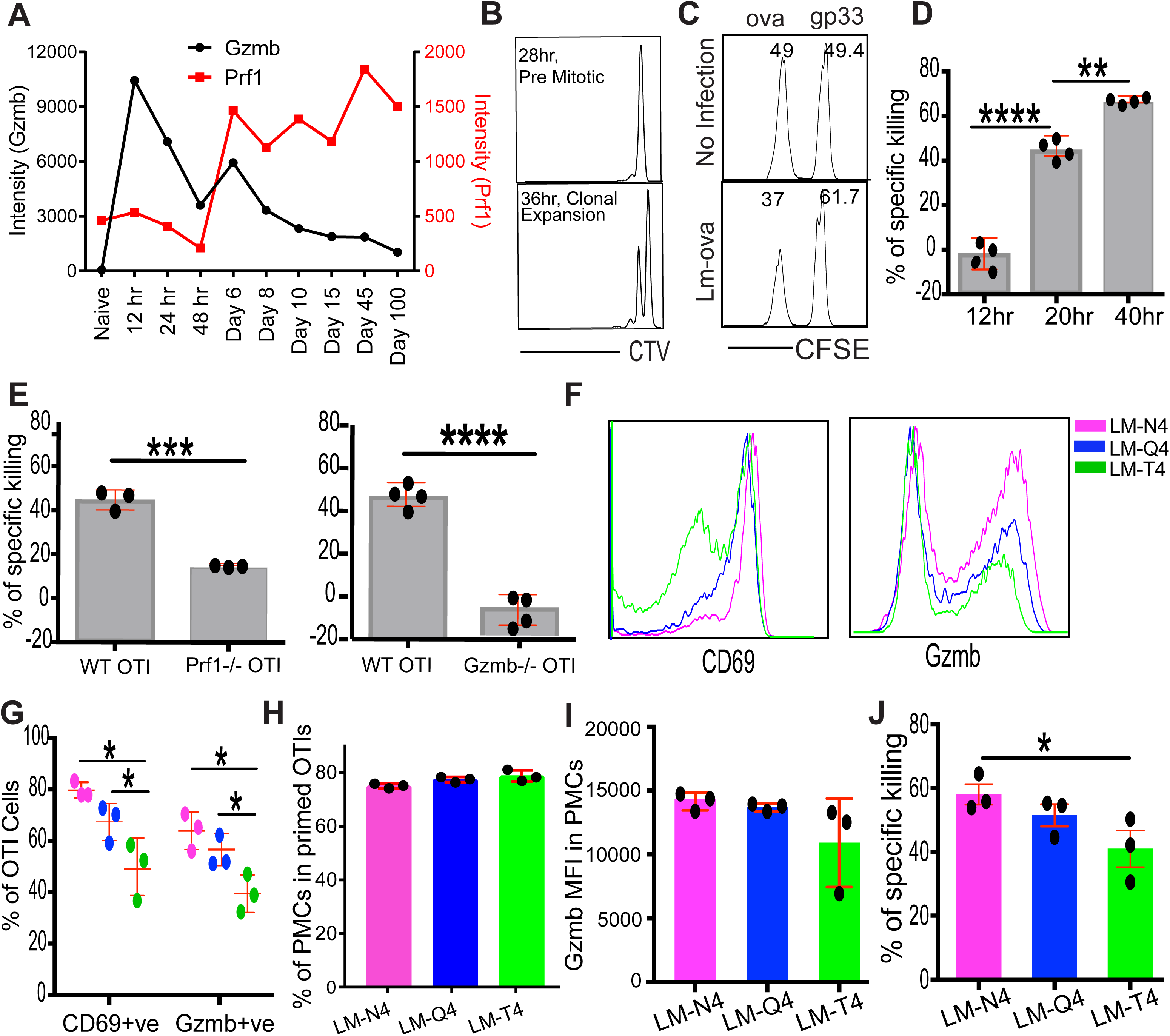
Pre-mitotic cytotoxicity during priming of OTIs in vivo. B) Temporal mRNA expression profiles of cytotoxic molecules in adoptively transferred OTI CD8 T cells in response to *Lm-ova* infection. The data shown is from the Immgen consortium (https://www.immgen.org). For the 12, 24, and 48 hour time-points, OTIs were transferred into mice after they were infected with *Lm-ova* for 24 hours. We call this ‘post-infection transfer’ model. This modified approach was used to synchronize T cell activation among the OTIs and clearly define the duration of priming. B) OTIs are in the pre-mitotic phase at least for the first 28 hours of priming after post-infection transfer. By 36 hours, some primed cells undergo their first division based on dilution of Cell Trace Violet (CTV) in divided cells. C) Antigen-specific, in vivo cytotoxic potential of primed OTIs assessed 20 hours after post-infection transfer. Killing of differentially labelled, antigen-specific target cells (ova-loaded) in the infected mouse can be deduced based on their reduced numbers compared to the nonspecific (gp33-loaded) targets. Relative numbers of antigen-specific and non-specific targets from the non-infected and infected mouse is used to calculate the % of specific killing. D) Quantitative comparison of specific killing activity assessed after varying durations following post-infection transfer. E) In vivo cytotoxicity of PMCs is dependent on granzyme B and a minor fraction of splenocyte targets can be killed by PMCs in a perforin-independent manner, as shown by using *Gzmb -/-* and *Prf1 -/-* OTIs respectively. Here in, OTIs were transferred 24 hours post-infection along with the targets and the targets were analyzed 20 hours later. Negative specific killing by *Gzmb -/-* OTIs means that ova-loaded targets were more than gp33-loaded targets on a relative basis in those samples. F) Histograms for CD69 and granzyme B probes are shown for OTIs stimulated by *L. monocytogenes* expressing variants of ovalbumin that give rise to APLs of SIINFEKL at the 4^th^ residue. Staining was performed 24 hours after post-infection transfer. G) Decrease in the percentage of OTIs that express CD69 and granzyme B with decreasing antigenic potency. H) Granzyme B is expressed in essentially the same % of primed OTIs (CD69+ve) irrespective of antigenic potency. I) Comparable amount of granzyme B is expressed in PMCs, irrespective of antigenic potency. J) Specific killing, measured 20 hours after post-infection transfer, decreases with antigenic potency. However the decrease is less dramatic than the decrease in the potency or affinity of the ova APL. Each data point in the figure represents a separate mouse. The bars represent mean values. Standard deviation is shown in red. Statistical significance of difference in values was calculated by paired t test. p values from two-tailed tests are denoted as follows in the figures: *p < 0.05, **p < 0.01, ***p < 0.001, ****p < 0.0001.

We assessed the dependency of PMC mediated killing of splenic targets on perforin and granzyme B. For this purpose, we used perforin-deficient (*Prf1-/-*) and granzyme B-deficient (*Gzmb -/-*) OTIs in *in vivo* killing assays at the 20 h time point. While there was significant pre-mitotic cytotoxicity in the absence of perforin, the specific targets were slightly protected in the absence of granzyme B, perhaps due to displacement of small numbers of wild-type endogenous PMCs from targets by granzyme B-deficient OTI PMCs (Figure 1E). Thus, cytotoxicity of PMCs is fully dependent on granzyme B, and partly dependent on perforin when whole splenocytes are used as targets *in vivo*. While *Prf1* mRNA was not increased at 12-48 hours, there is a significant basal level of *Prf1* mRNA that only slightly decreases from 12-48 hours (Figure 1A). Thus, there is some perforin activity that contributes to PMC mediated killing of splenic targets.

We next determined the role of potency of the antigen in the induction of PMCs. To assess this, we took advantage of *Lm-ova* variants that give rise to pMHC of known potency against the OTI TCR (Zehn et al., 2009). The natural ligand SIINFEKL (N4) has the highest potency; SIIQFEKL (Q4) has intermediate potency (39-fold lower based on dose); and SIITFEKL (T4) has the lowest potency (122-fold lower) (Daniels et al., 2006). Despite such large decreases in potency, both Q4 and T4 induced significant, although decreased, number of OTIs to express granzyme B at 24 hours after post-infection transfer (Figure 1F and 1G). In fact, among the primed (CD69^+^) cells, essentially the same % of cells express granzyme B, implying that development of PMCs among primed cells is independent of antigen potency (Figure 1H). Moreover, the expression levels of granzyme B on a per cell basis were not significantly different across the range of potencies (Figure 1I). Accordingly, specific killing activity mirrors the fraction of granzyme B expressing cells within the transferred OTIs (Figure 1J). Thus, PMCs develop efficiently in response to pMHC across a wide range of potencies.

### Extent of activation dictates the development of PMCs among primed cells

Strict correlation between the status of granzyme B expression and specific killing activity (Figure 1G and 1J), implies that granzyme B expression is the critical determinant for the development of PMCs in primed OTIs. We asked which of the primed CD8 T cells express granzyme B and thereby give rise to PMCs. We considered three transcription factors as putative regulators of granzyme B expression in the primed CD8 T cells. T-bet is a known activator of granzyme B expression in the effector phase (Sullivan et al., 2003). IRF4 and cMyc are two of the master regulators of the priming phase and initial clonal expansion (Mayya and Dustin, 2016). As expected, expression of the 3 transcription factors, after 24 hours of priming, was dictated by the potency of the pMHC (Figure 2A). When quantified, IRF4 showed the best scaling among the considered transcription factors. Expression of IRF4 is known to be scaled as per the potency and dose of pMHC and the duration of TCR signalling (Nayar et al., 2014). Expression level of IRF4 then dictates the scaled expression of proteins driving energy production, anabolic processes and proliferation (Man et al., 2013). CD69^+^ OTIs cells that expressed granzyme B protein had higher expression of the transcription factors compared to those that did not express granzyme B (Figure 2B, top). Among the considered transcription factors, IRF4 showed the highest increase in expression within PMCs relative to those primed OTIs that did not express granzyme B (Figure 2B, right). Similarly, expression of IRF4 correlated the most with granzyme B expression when cell-to-cell variability was considered among the PMCs (Figure 2B, left). Cells that expressed the highest levels of IRF4 and Myc were expected to be larger owing to increased anabolic metabolism. Accordingly, we found that cells with higher forward (related to volume) and side (related to granularity) scatter 24 hours after infection were more enriched for granzyme B expression (Figure 2C). Conversely, PMCs had higher forward and side scatter (Figure 2D). Thus, the extent of activation is a major determinant of granzyme B expression and thereby the development of PMCs.

**Figure 2:**
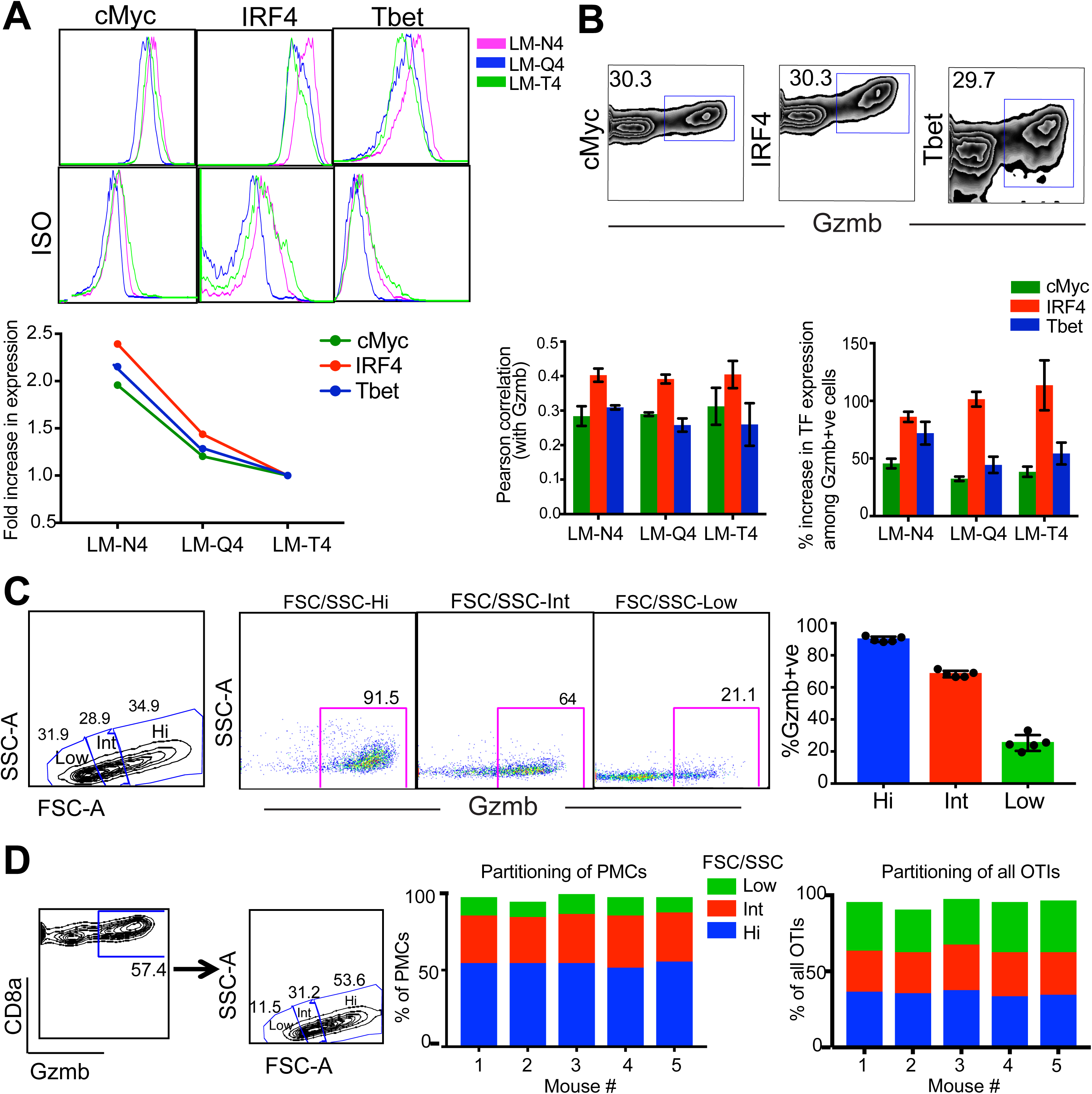
Extent of activation dictates the development of PMCs. A) Increase in the expression of transcription factors, namely cMyc, IRF4 and Tbet, during the priming phase with *Lm-ova* with increasing antigenic potency. OTIs were transferred post-infection and stained 24 hours later for the transcription factors. Geometric mean fluorescence of the population is considered to look at fold-increase in the expression of the transcription factor. Average from 3 mice are shown. B) Relationship between the expression of the transcription factors and that of granzyme B among OTIs that are CD69+ve in response to *Lm-ova* that expresses the N4 APL. Pearson correlation was calculated for cells that express granzyme B. High degree of correlation, especially with IRF4, suggests that highly activated cells express granzyme B. Data from 3 mice are shown with the bar height representing mean value and error bars representing standard deviation. C) OTIs were grouped into roughly equal sized three bins, namely Low, Int, and Hi, with increasing forward and side scatter. Extent of enrichment of PMCs (i.e. % of Gzmb+ve OTIs) in the three bins increased with increase in forward and side scatter, which are proxies for anabolic activity of T cells. D) PMCs were placed into the same bins as above to assess how they partition into the three bins. Majority of PMCs fell in the bin with highest forward and side scatter, followed by the bin with intermediate forward and side scatter. Roughly equal partitioning of all OTIs into the three bins is also shown for comparison. The subpopulations do not add up to 100% as some of the cells were outside of all three bins.

### Comparison of cytotoxicity of PMCs and conventional effector CTLs

In our effort to further characterize the killing mechanism operating in PMCs we compared them with CTLs using an ex vivo cytotoxicity assay, which provided us with superior control of effector:target (E:T) ratios and equivalent settings for cells that may not co-exist in vivo. For CTLs, we performed a standard adoptive transfer of 10^3^ OTIs, infected with 5000 *Lm-ova* after 24 hrs, waited an additional 5 days and sorted using an allotypic marker. For PMCs, we sorted OTIs using the same markers after 20 hours following post-infection transfer. Targets with cognate and control peptides were labelled with different levels of CFSE in exactly the same manner as for *in vivo* cytotoxicity assay, except that the cells interacted in co-culture for 12 hours prior to analysis by flow cytometry (Figure 3A, left). At E:T ratio of 0.5:1, the CTLs appeared to be 2-fold more potent than the PMCs (Figure 3A, middle). However, titration of the E:T ratio revealed saturation in the % of splenic target killing: at ~80% for CTLs and at ~50% for PMCs (Figure 3A, right). This suggests that the main advantage of CTLs over PMCs is the ability to kill a larger range of targets within the heterogeneous splenocyte population.

**Figure 3:**
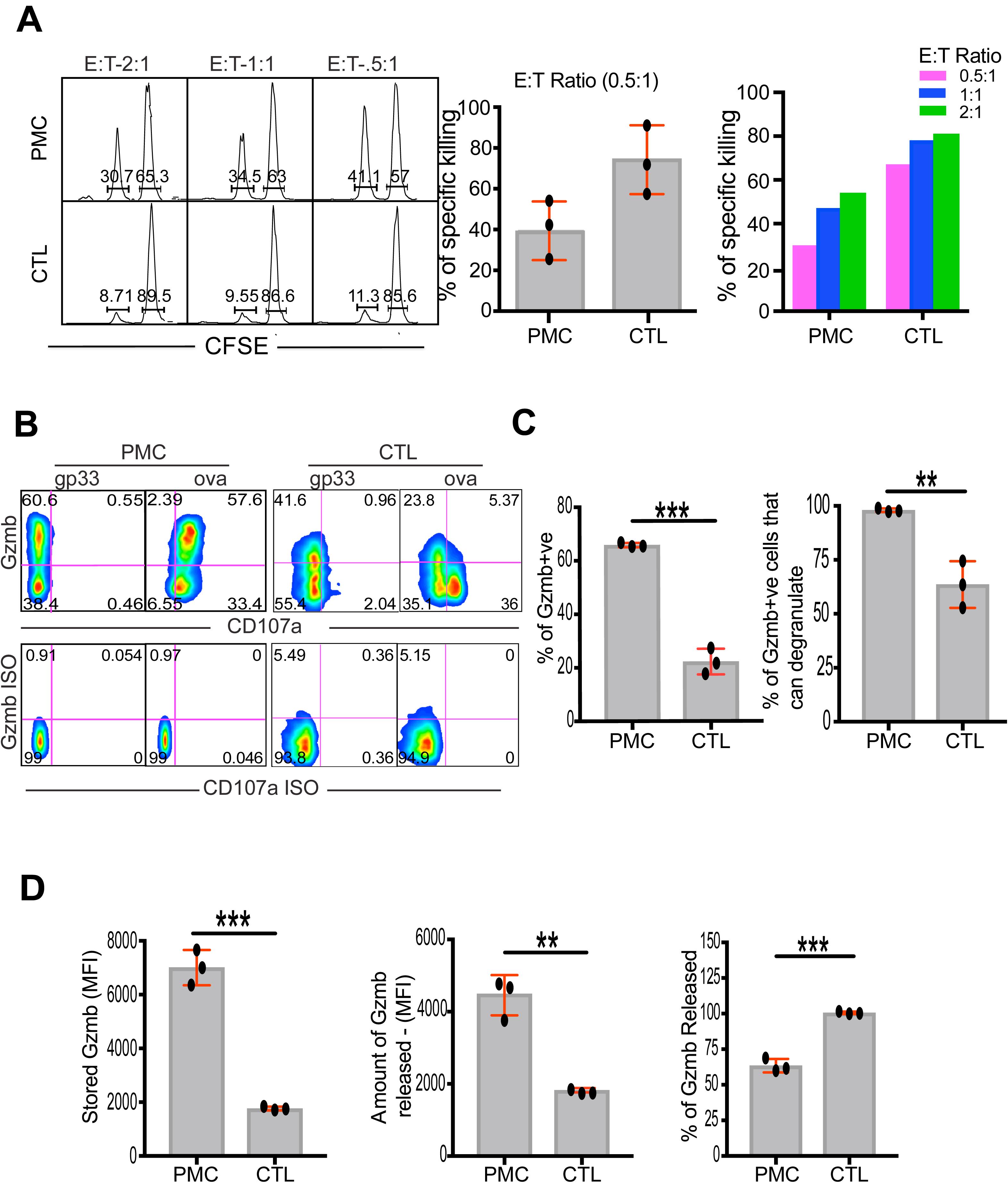
Ex vivo comparison between PMCs and conventional effector CTLs that arise in response to *Lm-ova* infection. Comparisons between PMCs isolated 20 hours after post-infection transfer and conventional effectors (CTLs) isolated 5 days after post-infection transfer of naïve OTIs. A) Specific killing was measured after 12hrs of co-culture at three different effector-to-target (E:T) ratios. Antigen-specific target (ova-loaded) cells are distinguishable from irrelevant targets (gp33- loaded) by reduced CFSE labelling in the histogram plots. Specific killing activity of CTLs is ~2-fold higher than that of PMCs. B) Dual staining of intracellular granzyme B and surface CD107a, the degranulation marker, in PMCs and CTLs. PMCs or CTLs were co-cultured with target cells for 1 hour before staining. Quantitative comparison between PMCs and CTLs stimulated with ova-loaded vs. gp33-loaded targets revealed several aspects of cytotoxicity. C) Estimation of fraction of bona fide killers based on dual staining experiments shown in B. % of Gzmb+ve cells is defined based on isotype staining. Gzmb+ve cells that can degranulate defines the subset of *bona fide* killers. It is calculated using the following formula:

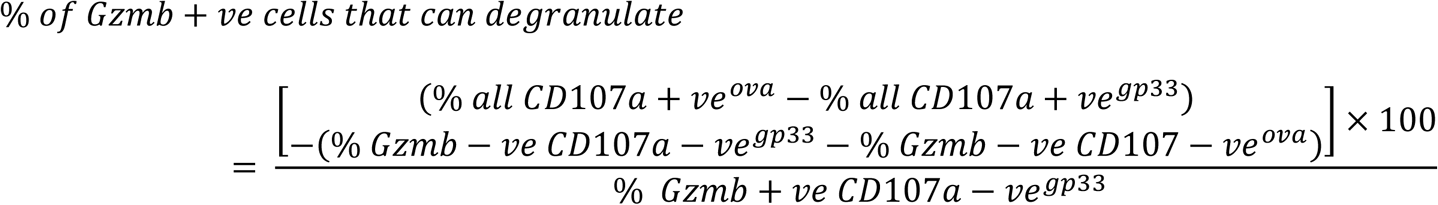 Based on this, 65.8% × 98% = 64.5 % of cells in the PMC population are bona fide killers and 22.2% × 63.6% = 14.1% of cells in the effector CTL population are bona fide killers. D) Amount of stored granzyme B is calculated based on geometric mean fluorescence intensity in the cells engaged with gp33-loaded target cells. Amount of granzyme B released is calculated using the following formula:

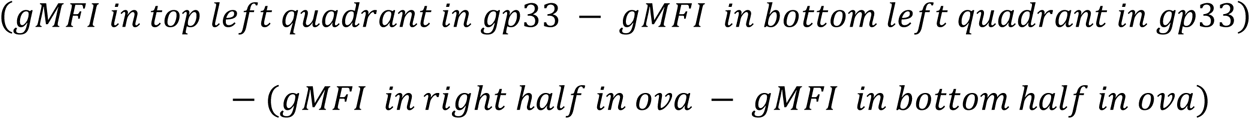 PMCs have a lot more granzyme B in storage than CTLs, however they release only a fraction of it by degranulation in an hour. Each data point in the figure represents a separate mouse. The bars represent mean values. Standard deviation is shown in red. Standard deviation is shown in red. Statistical significance of difference in values was calculated by paired t test. p values from two-tailed tests are denoted as follows in the figures: *p < 0.05, **p < 0.01, ***p < 0.001, ****p < 0.0001.

Rapid killing of targets by CTLs requires exocytosis (or degranulation) of perforin and granzyme B from secretory lysosomes, which exposes the lysosomal membrane protein CD107a transiently on the cell surface (Stinchcombe and Griffiths, 2007). In order to understand how the killing potency manifests in the two differentiation stages, we performed dual staining of intracellular granzyme B and cell surface CD107a after an hour of co-culture with splenic target cells (Figure 3B). All processing was performed at the same time and instrument settings were kept constant to allow comparisons between PMCs and CTLs. Both PMCs and CTLs displayed antigen-dependent exposure of CD107a on the surface of most cells regardless of intracellular granzyme B (Figure 3B). Quantification revealed that, on a relative basis, only about 1/3^rd^ of conventional CTLs express granzyme B and 2/3^rd^ among those are actually able to degranulate their stored granzyme B (Figure 3C, graphs show absolute comparisons). Thus, the number of bona fide killers among CTLs is only ~22% of that of PMCs. With ~2-fold higher specific killing, this would mean that killing potency of effector CTLs is ~9-fold higher than that of PMCs for splenic targets. The effector CTLs have about 3.5-fold less granzyme B in storage than PMCs (Figure 3D, left), which is consistent with mRNA levels (Figure 1A). The PMCs display relatively less antigen driven depletion of granzyme B during the assay (Figure 3D, right). Due to higher absolute expression levels, PMCs appear to release over 2-fold more granzyme B per cell than conventional effectors, assuming no antigen-dependent replenishment during the one hour of the assay in both stages (Figure 3D, middle). In summary, PMCs share with CTLs the ability to degranulate and release granzyme B in an antigen specific manner. But PMCs are considerably less potent and appear unable to kill some splenic targets that are killed by CTLs.

### PMCs kill conventional DCs in a granzyme-B dependent manner

After establishing that *L. monocytogenes* and LCMV infection generate PMCs, we set out to examine their physiological role. Imaging of antigen specific CD8 T cells interacting with *L. monocytogenes* infected DCs in the splenic white pulp indicate that durable interactions are established as soon as a naïve CD8 T cell encounters an infected DC (Aoshi et al., 2008). Primed CD8 T cells are found interacting with DCs even 38 hours after infection with vaccinia virus (Eickhoff et al., 2015). We used immunofluorescence staining of fixed tissue from CD11c-YFP mice to investigate the expression of granzyme B and interaction of transferred OTIs and endogenous CD8 T cells with DCs in the splenic white pulp during the pre-mitotic phase. We transferred 10^6^ OTIs post-infection to facilitate imaging of T cell-DC interaction after up to 20 and 36 hours of priming, respectively, for OTI and endogenous T cells. We observed OTIs and endogenous CD8 T cells expressing granzyme B in the same tissue sections (Figure 4A). Most of these cells had punctate distribution of granzyme B consistent with detection of granules (Figure 4A). Some of these cells also shared a large interface with DCs, suggesting immune synapse formation (Figure 4A, bottom panels). Upon manual enumeration, we found ~140 OTIs per mm^2^ of the tissue sections, of which ~40 per mm^2^ showed visibly distinct granzyme B staining and ~30 OTIs per mm^2^ (~20% of all OTIs) were present as conjugates with DCs (Figure 4B). Similarly, we found ~360 endogenous CD8 T cells per mm^2^ that were expressing granzyme B, which we estimate to be about 2% of all endogenous CD8 T cells in the tissue sections (Figure 4C). In comparison with OTIs primed for 20 hours, we saw far less endogenous PMCs that were also interacting via large interface with DCs (Figure 4C). This is consistent with disengagement of primed T cells from DCs just prior to the commencement of division, i.e. the end of pre-mitotic phase (Bohineust et al., 2018). We further confirmed the presence of endogenous PMCs by flow cytometry (Figure 4D). Based on intracellular staining in isolated cells, we enumerated ~5×10^4^ endogenous PMCs in the spleen 24 hours after infection (Figure 4D).

**Figure 4:**
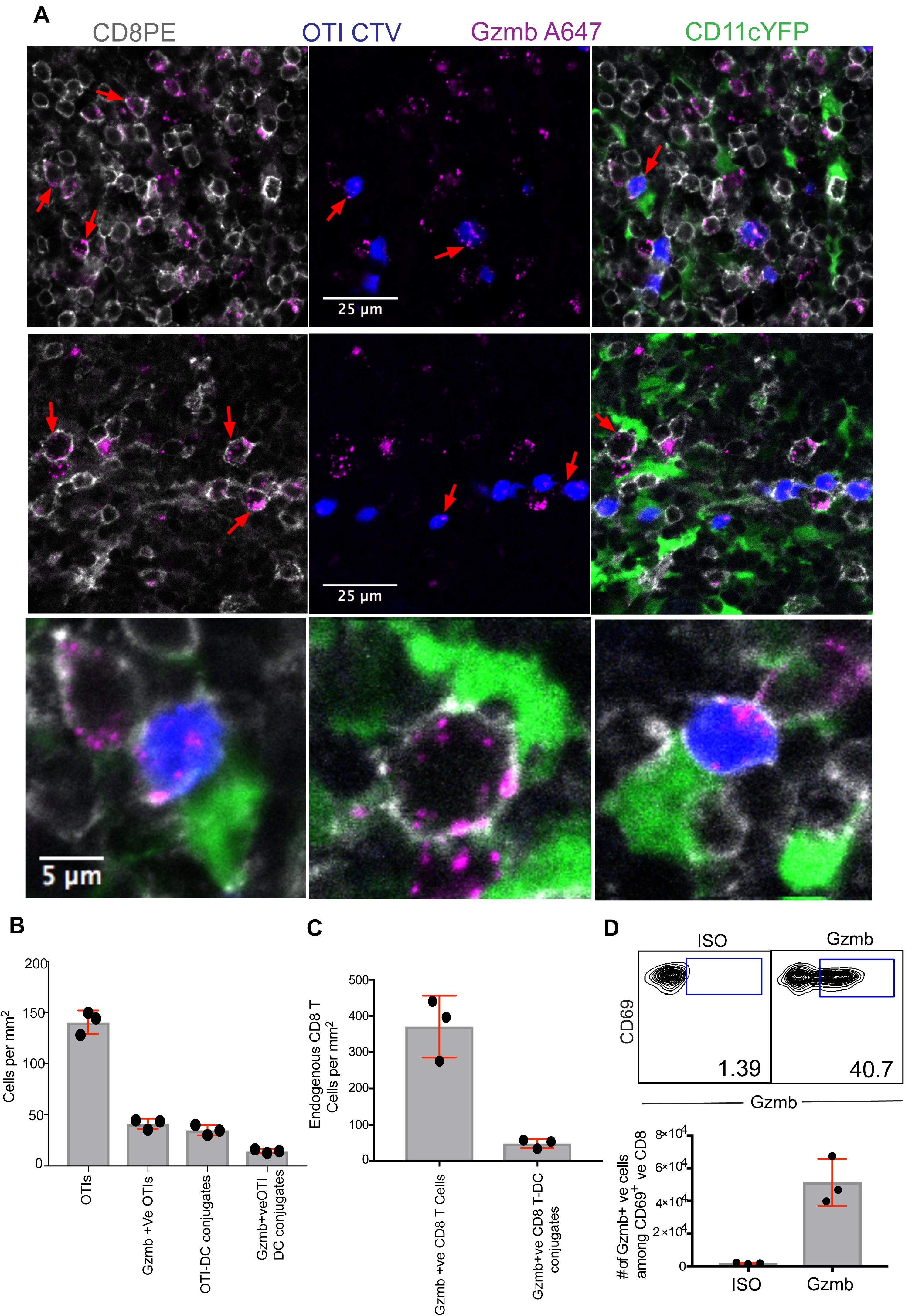
OTI and endogenous PMCs engage with DCs in situ. A) Immunofluorescence images of endogenous and OTI PMCs in T cell-rich areas of the white pulp. We had pre-established that all the transferred cells are located in the white pulp 20 hours after transfer, based on IgD staining to identify B cell follicles in the white pulp. OTIs were labelled with CTV (shown in blue) and transferred 24 hours post-infection. DCs were identified by transgenic YFP expression (shown in green). Endogenous CD8 T cells were identified by staining with a phycoerythrin-conjugated CD8 antibody (shown in grey). OTIs in these tissue sections have undergone 20 hours of priming whereas endogenous CD8 T cells have undergone 36 hours of priming. The top two rows are images of two different sections with overlays of different channels shown in the same row. Punctate distribution of granzyme B was observed in both endogenous and OTI PMCs, as highlighted by red arrows in the left and centre panels. Some of the T cells were also found to be closely interacting with DCs, as highlighted by red arrows in the right panels. These examples have been magnified in the bottom row for better appreciation of features. OTIs (in B) and endogenous T cells (in C) were counted from many fields of 270µm size giving a total sampling area of 1.1mm^2^ and 0.75mm^2^, respectively. Only visibly distinct granzyme B staining was considered while manually counting granzyme B+ve cells. For T-DC conjugates, the T cells with >20% of coverage of the perimeter by the interface were manually identified. D) Percentage (top) and absolute number (bottom) of endogenous PMCs quantified by staining for intracellular granzyme B and flow cytometry 24 hours after infection with *Lm-ova*. Each data point in the figure represents a separate mouse. The bars represent mean values. Standard deviation is shown in red.

Immunofluorescence imaging suggested that DCs themselves were the likely targets of PMCs during priming. As suggestive evidence, we observed increasing amount of granzyme B in CD11c^+^ DCs during the pre-mitotic phase, presumably due to release by the PMCs and uptake by the DCs (Figure S2A). If this were to be true and of functional relevance, the number of viable DCs in the spleen would be reduced after infection with *L. monocytogenes*. Accordingly, we observed that the number of CD4^+^ and Dec205^+^ DCs decreased drastically within the first 24 hours and over the next 24 hours these populations almost become undetectable (Figure 5A). In contrast, the number of CD8α^+^ DCs remained similar over the first 24 hours and then reduced by about 60% over the next 24 hours. Both these observations are consistent with a previous report (Mitchell et al., 2011). Similarly, we observed drastic reduction in the number of CD4^+^ and Dec205^+^ DCs, but not in CD8α^+^ DCs, 48 hours after infection with LCMV (Figure S2C). The refractory nature of CD8α^+^ DCs may be due to significantly higher expression of Spi6 (also called PI9) in this subset of DCs (Figure S2B), which is an inhibitor of granzyme B (Lovo et al., 2012). We further confirmed that the reduction in the number of DCs was dependent on PMCs, as there was no change in the number of DCs when mice deficient in CD8 T cells (CD8α -/- mice) were infected (Figure 5B). Isolated PMCs were also capable of killing DCs ex vivo, in an antigen-specific manner (Figure 5C). We observed ~45% specific-killing of enriched CD11c^+^ DCs after 12-16 hours of co-culture with PMCs ex vivo. This provides further evidence that reduction observed in vivo was in fact due to killing by PMCs and not due to other phenomena.

**Figure 5:**
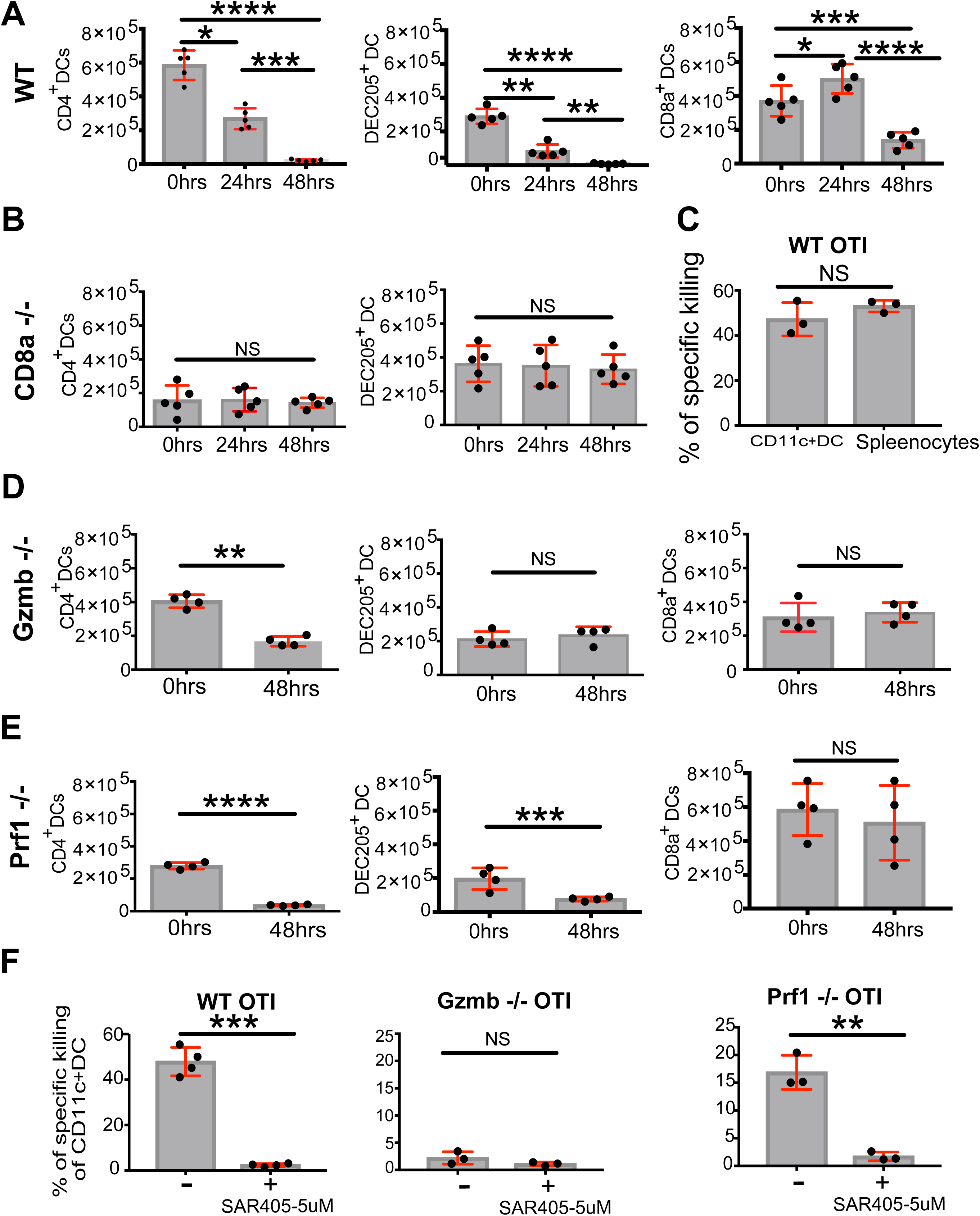
PMCs kill conventional DCs in a granzyme-B dependent and largely perforin-independent manner. Absolute counts of conventional DC subsets in the spleen of wild-type B6 (in A), CD8α -/- (in B), *Gzmb -/-* (in D), and *Prf1 -/-* (in E) mice before infection (labelled 0 hrs) and 24 and 48 hours after infection with *Lm-ova*. A) More than 95% of CD4+ and Dec205+ DCs disappear in the first 48 hours in wild-type B6 mice. In contrast, the decrease is ~60% in the CD8α+ DC subset. B) No decrease in the number of DCs in CD8α -/- mice after infection. Reduced number of CD4+ DCs in CD8α -/- mice compared to the B6 mice is likely because of reduced T-cell area niche in the white pulp resulting from lack of CD8 cells. C) Isolated OTI PMCs efficiently kill CD11c+ DCs ex-vivo. Enriched DCs were loaded with ova or gp33 and co-cultured with isolated PMCs for 12-16 hours before measuring % of specific killing of DCs. D) No reduction in the number of Dec205+ and CD8α+ DCs and ~60% reduction in that of CD4+ DCs in *Gzmb -/-* mice after infection. E) Drastic reduction in the number of CD4+ and Dec205+ DCs in *Prf1 -/-* mice after infection. F) Inhibition of ex vivo killing of CD11c+ DCs by the Vps34 inhibitor SAR405 (left). Ex vivo killing of DCs is dependent on granzyme B (middle). Perforin deficient OTI PMCs can kill some of DCs ex vivo (left). Each data point in the figure represents a separate mouse. The bars represent mean values. Standard deviation is shown in red. Standard deviation is shown in red. Statistical significance of difference in values was calculated by paired t test. p values from two-tailed tests are denoted as follows in the figures: *p < 0.05, **p < 0.01, ***p < 0.001, ****p < 0.0001.

Granzyme B is the only cytotoxic molecule that PMCs express in an inducible manner, with perforin expression remaining constitutive and low (Figure 1A and Figure S1A). Therefore we asked if killing of splenic DCs by PMCs and the resultant dramatic reduction in number of viable DCs within two days of infection is granzyme B dependent. There was no reduction in the number of Dec205^+^ and CD8α^+^ DCs within 48 hours of infecting *Gzmb -/-* mice (Figure 5D). Even in the more sensitive CD4^+^ DC population, there was ~60% pruning in *Gzmb -/-* mice, whereas >95% reduction were lost in wild-type mice (Figure 5D). Thus, killing of splenic DCs by PMCs was dependent on granzyme B. Residual killing of CD4+ DCs in *Gzmb -/-* mice is likely because of granzyme C, which can cause cell death without activating caspases (Johnson et al., 2003). It is known that CTLs from *Gzmb -/-* mice express more granzyme C (Cai et al., 2009). Expression of granzyme C was induced in the pre-mitotic phase in the wild-type setting (Figure S2D), and the chromosomal deletion in the *Gzmb -/-* mice may facilitate the expression of granzyme C to reach efficacious levels. It is also possible that intracellular infection, necroptosis and pyroptosis contribute to death of DCs in wild-type and *Gzmb -/-* mice (Carrero et al., 2004; Man and Kanneganti, 2016; Vandenabeele et al., 2010), but the retention of DC populations in CD8α -/- mice make these explanations less likely.

Expression of perforin is considerably lower in PMCs (Figure 1A). Therefore, we asked if the killing of DCs in vivo is independent of perforin. There was drastic decrease in numbers of CD4^+^ and Dec205^+^ DCs within two days of infecting *Prf1 -/-* mice (Figure 5E). Thus, killing of CD4^+^ and Dec205^+^ DCs is essentially perforin-independent. Perforin can oligomerise into a channel on the plasma membrane allowing direct entry of released granzyme B into the target cytosol (Voskoboinik et al., 2010). Released granzyme B can also bind to the plasma membrane and then be taken up by endocytosis or pinocytosis (Bird et al., 2005; Shi et al., 2005). DCs are equipped with multiple mechanisms such as endocytosis, pinocytosis, and phagocytosis, to take up material from the extra cellular milieu (Trombetta and Mellman, 2005). We reasoned that uptake of granzyme B is very efficient in DCs and asked if preventing the uptake impacts killing of DCs. We chose to target Vps34, a highly conserved class III phosphatidylinositol 3-Kinase that is essential for membrane and vesicle trafficking and thus is involved in antigen uptake pathways, autophagy and cross-presentation (Backer, 2008; Parekh et al., 2017). Inhibition of Vps34 using 5µM of SAR405 (Ronan et al., 2014) prevented ex vivo killing of CD11c^+^ DCs by PMCs (Figure 5F, left). Consistent with the in vivo data, *Gzmb -/-* PMCs do not kill DCs ex vivo (Figure 5F, middle) and PMCs lacking perforin kill a substantial number of CD11c^+^ DCs. However, even perforin-independent killing is abrogated when Vps34 is inhibited (Figure 5F, right). These results highlight the importance of antigen uptake pathways in mediating killing of splenic DCs by PMCs.

### Pre-mitotic cytotoxicity terminates ongoing antigen presentation and prevents recruitment of additional antigen-specific naïve T cells

We probed if and when elimination of splenic DCs by PMCs leads to decrease and termination of ongoing antigen presentation. Transferred, antigen-specific T cells can be used as reporters of ongoing and subsequent antigen presentation by quantifying the extent of their proliferation (Hafalla et al., 2003; Wong and Pamer, 2003). There was considerable proliferation of OTIs transferred a day after infection with *Lm-ova*. However, proliferation in cells transferred later on dropped, with virtually no antigen presentation 3 days after infection to support activation and initiate division of OTIs transferred at that time (Figure 6A). Thus, antigen presentation is essentially terminated a day after nearly all the CD4^+^ and Dec205^+^ splenic DCs and nearly ~60% of CD8^+^ splenic DCs are eliminated (Figure 5A). Next, we asked if termination of antigen-presentation is also dependent on granzyme B, like the killing of splenic DCs. We used P14 TCR transgenic mice as hosts as they are not expected to give rise to any endogenous PMCs when infected with *Lm-ova* (Figure 6B). We then transferred wild-type, or *Gzmb -/-* or *Prf1 -/-* OTIs to give rise to PMCs in the P14 host upon infection. With the wild-type OTIs, antigen presentation was essentially terminated within 2 days, as reporter cells (CTV labelled OTIs that carry a congenic marker) do not show any proliferation over the next 3 days. However, with *Gzmb -/-* OTIs, antigen presentation continues beyond 2 days, as the reporter cells proliferated. In contrast, with *Prf1 -/-* OTIs antigen presentation is considerably diminished after 2 days (Figure 6B). Thus, termination of antigen presentation by PMCs is also dependent on granzyme B but largely independent of perforin. As with residual killing of splenic DCs in *Gzmb -/-* mice, the extent of antigen presentation is reduced by *Gzmb -/-* OTIs. This inference is made based on larger extent of proliferation observed in reporter cells in infected P14 mice when pre-mitotic cytotoxicity was totally absent without the initial transfer of OTIs. Thus, results from these antigen presentation assays mirror the extent of decrease seen in numbers of conventional splenic DCs after infection, both in wild-type mice and in other genetic backgrounds examined. These results also prove that pre-mitotic cytotoxicity prevents recruitment of additional, ‘late-comer’ antigen-specific T cells to the response (D’Souza and Hedrick, 2006).

**Figure 6:**
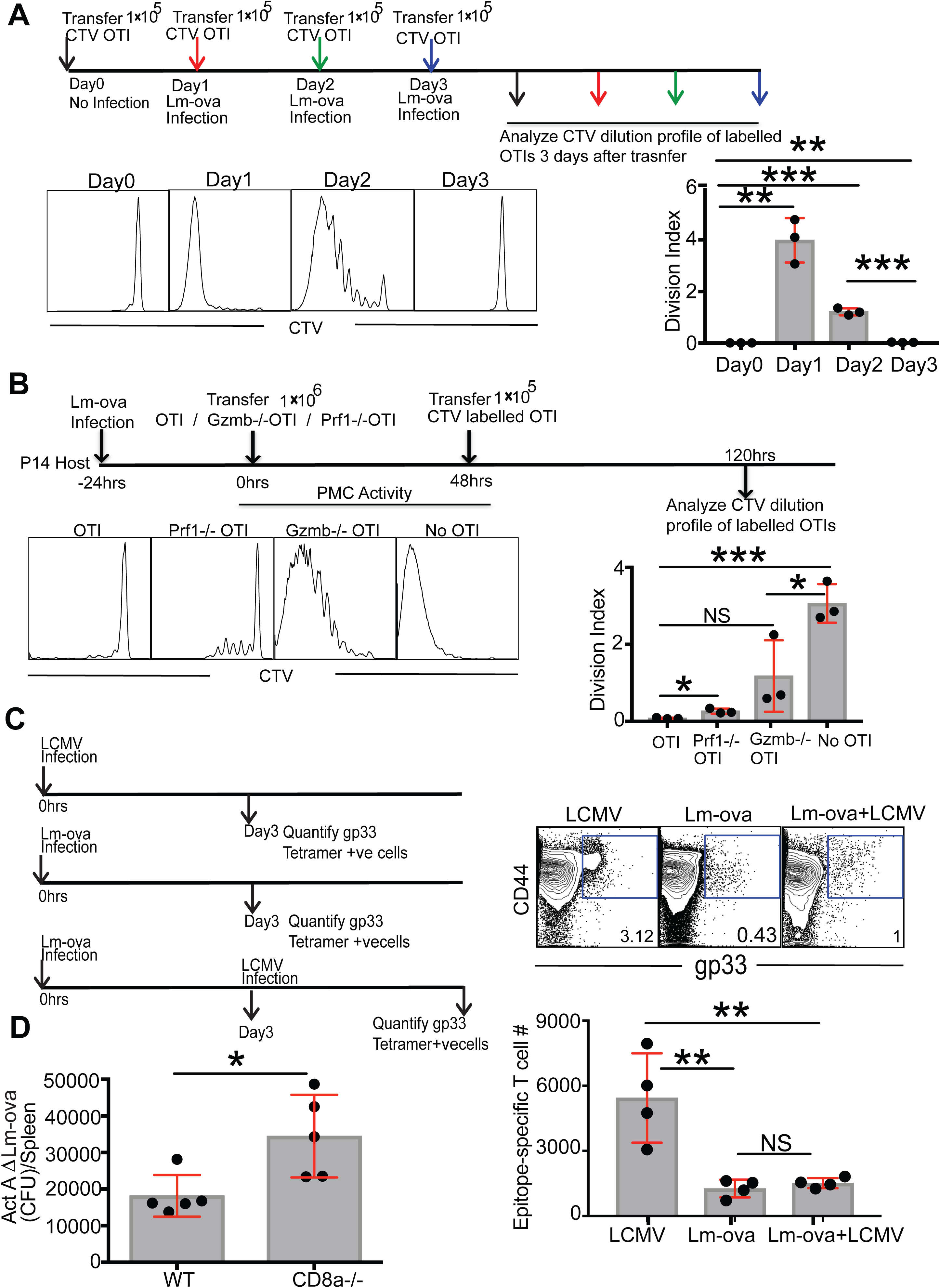
Immunological consequences of elimination of conventional DCs by PMCs. A) Reduction and termination of antigen presentation after *Lm-ova* infection. The assay schematic is shown on top. CTV labelled, congenic reporter OTIs were transferred on every subsequent day following infection. Proliferation of the reporter cells is dependent on the extent and duration of antigen presentation that they experience. Representative CTV dilution histograms of reporter OTIs are shown. The extent of proliferation 3 days after transfer is quantified using ‘Division Index’ that is based on division peaks identified in the CTV dilution histogram using the FlowJo software. Division index is the average number of divisions that a cell has undergone in the original reporter population. B) Termination of antigen presentation is dependent on granzyme B but largely independent of perforin. Wild-type OTIs or *Gzmb -/-* or *Prf1 -/-* OTIs were transferred into P14 hosts following infection. The transferred OTIs are, presumably, solely responsible for pre-mitotic cytotoxicity and killing of DCs in the P14 hosts. Residual antigen-presentation is assayed with congenic reporter OTIs as above in A. There was considerable proliferation of reporter cells in P14 hosts that had received *Prf1 -/-* OTIs but not in hosts that had received *Gzmb -/-* OTIs. C) Suppression of response to LCMV infection 3 days after infection with *Lm-ova*. The experiment schematic is shown on left. Endogenous gp33-specific T cells were identified using tetramer staining in enriched CD8s from splenocytes 3 days after infection. The number of gp33-specific CD44+ve CD8 T cells did not increase upon LCMV infection if they had previously been infected with *Lm-ova* due to lack of proliferation. D) Increased burden of ActA deficient *Lm-ova* in CD8α -/- mice that lack CD8 T cells. The mice were infected with 5×10^4^ CFU ActA deficient *Lm-ova* and the number of expanding bacteria 48 hours later in the spleen were counted by growing serial dilutions of splenic tissue homogenate on agar plates. Each data point in the figure represents a separate mouse. The bars represent mean values. Standard deviation is shown in red. Standard deviation is shown in red. Statistical significance of difference in values was calculated by paired t test. p values from two-tailed tests are denoted as follows in the figures: *p < 0.05, **p < 0.01, ***p < 0.001, ****p < 0.0001.

### Pre-mitotic cytotoxicity undermines response to subsequent secondary infection

Rapid elimination of DCs prevents activation of reporter T cells specific to the primary infection as they were introduced after the termination of antigen presentation. This implies that additional antigen-specific naïve T cells entering the lymphoid organ are not recruited to the response. In addition, termination of antigen presentation also implies that the host will be hampered in mounting a T cell response against a different pathogen a few days after the initial infection. To test if this is indeed the case, we infected mice with LCMV 3 days after the initial infection with *Lm-ova* (Figure 6C). We found that, endogenous gp33-specific CD8 T cells did not undergo any clonal expansion over 3 days following the introduction of LCMV as a second pathogen. Thus, pre-mitotic cytotoxicity in response to the primary infection can profoundly impair adaptive immunity to an immediate secondary infection.

### Pre-mitotic cytotoxicity limits the spread of DC-borne infections

Several intracellular pathogens, be it viruses, bacteria or protozoans, can directly infect DCs (Table S1). In a number of these cases, it is either known or suspected that the pathogens use DCs for further dissemination and/or establishment of infection (Table S1). Rapid elimination of the DCs by PMCs could aid in controlling the overall pathogen burden. We set out to test that hypothesis in the case of *L. monocytogenes* as DCs are critical permissive carriers of *L. monocytogenes* for dissemination. *L. monocytogenes* in the blood stream are captured at the marginal zone in the spleen by macrophages and DCs. Infected CD8α+ DCs then move into T-cell rich areas of the white pulp due to maturation elicited by innate sensing (Aoshi et al., 2008). Thereby, *L. monocytogenes* use CD8α^+^ DCs to infiltrate the white pulp, expand and then spread rapidly into lymphocytes (Edelson et al., 2011; Neuenhahn et al., 2006). However, the intercellular dissemination into lymphocytes and exponential growth within 24 hours occurs even before PMCs have a chance to act. Cell-to-cell spreading is absent in ActA deficient *L. monocytogenes*, thus reducing rapid dissemination and virulence (Kocks et al., 1992). We tested if pre-mitotic cytotoxicity can control the DC-borne spread of ActA deficient *L. monocytogenes*. CD8α -/- mice that do not give rise to PMCs harboured more expanding bacteria 48 hours after infection than wild-type B6 mice (Figure 6D). Thus, pre-mitotic cytotoxicity can control dissemination of the pathogen as long as it does not rapidly spread further from the DCs before pre-mitotic cytotoxicity emerges in the primed CD8 T cells.

### Pre-mitotic cytotoxicity limits clonal expansion in a self-regulatory fashion

Reducing the duration of antigen presentation has been shown to influence clonal expansion and differentiation of CD8 T cells (Blair et al., 2011). Herein, we have a scenario of extended duration of antigen presentation as a result of lack of or reduction in pre-mitotic cytotoxicity. First, we asked if availability of cytokines known to influence CD8 T cell expansion and differentiation is increased due to lack of pre-mitotic cytotoxicity. We did not see increased presence of IL12, IL6 and IL-1β two and three days post-infection despite continued presence of DCs in CD8α -/- mice (Figure S3A). In contrast, interstitial IL2 that is predominantly produced by activated CD4 cells, is significantly higher in splenic tissue homogenates from CD8α -/- mice and *Gzmb -/-* mice after infection (Figure 7A). This is presumably due to increased activation and/or increased numbers of responding CD4 cells. In order to test the effect of increase in the duration of antigen presentation on clonal expansion, we adoptively transferred 10,000 OTIs into wild-type, CD8α -/- and *Gzmb -/-* mice as reporter cells and then infected them with *L. monocytogenes* expressing a variant of ovalbumin that gives rise to T4 APL of the epitope SIINFEKL. As only ~1000 among these engraft, we surmised that PMCs arising out of these reporter cells will have a minor, insignificant contribution to endogenous pre-mitotic cytotoxicity, or lack thereof, that would determine the duration of antigen presentation. The choice of T4 APL in place of natural N4 epitope of ovalbumin was guided by the notion that lower strength of the TCR signalling would likely make duration of TCR signalling a major determinant of the extent of clonal expansion and differentiation. We observed a larger fraction of OTIs that had divided more than five times in the CD8α -/- mice three days after infection, implying higher rate of proliferation in a subset of progeny (Figure 7B). Accordingly, CD8α -/- mice that did not have any reduction in viable DCs after infection had ~7-fold more OTIs. High expression of CD25, the α chain of IL2 receptor, and prolonged availability of IL2 during the early part of clonal expansion phase has been shown to be predictive of the extent of further proliferation and of terminal differentiation in CD8 T cells (Kalia et al., 2010; Pipkin et al., 2010). Wild-type mice had the lowest percentage of CD25hi cells, followed by *Gzmb -/-* mice and then the CD8α -/- mice with the highest percentage of CD25hi cells among OTI reporter cells 3.5 days after infection (Figure 7C). Thus, the duration of antigen presentation dictates the expression of CD25. *Gzmb -/-* mice had ~2-fold more OTIs eight days after infection compared to the wild-type mice both in the spleen and liver (Figure 7D). Thus, prolonged antigen presentation in *Gzmb -/-* mice leads to increased expression of CD25 early during clonal expansion that eventually leads to increased rate of proliferation of responding CD8 T cells. Surprisingly, we did not observe any difference in the differentiation status of the effectors at this stage. Same fraction of SLECs and MPECs were found in both wild-type and *Gzmb -/-* mice (Figure S3B). We also examined the extent of clonal expansion of OTIs in response to *L. monocytogenes* expressing the natural N4 APL as a model high-affinity antigen. After 4.5 days of infection, there were ~2-fold more OTIs in the *Gzmb -/-* mice than in the wild-type B6 mice owing to higher rate of proliferation from prolonged antigen presentation (Figure 7E). However, 8 days after infection, the population of OTIs in *Gzmb -/-* mice had not expanded any further. This suggests that contraction of the OTI population started during the intervening period in the *Gzmb -/-* mice. In contrast, in the wild-type B6 mice the OTIs continued to have net proliferation (Figure 7E). As in the case of T4 APL, OTIs responding to the N4 APL gave rise to the same fraction of SLECs and MPECs in both wild-type and *Gzmb -/-* mice (Figure S3C, S3D). In each of the cases examined above, whenever the duration of antigen presentation was longer, the measured population size at different time-points implied higher rate of proliferation during the phase of clonal expansion. The peak size of the population is also influenced by the time of onset of contraction and rate of contraction. Duration of antigen presentation also appears to impact these features of population dynamics. Overall, we conclude that pre-mitotic cytotoxicity in primed CD8 T cells limits the rate of their own ensuing proliferation to reduce the number of effector cells produced during the expansion phase without impacting their differentiation trajectory. As majority of the PMCs arise out of highly activated T cells (Figure 2), pre-mitotic cytotoxicity is a negative feedback or an auto-inhibitory mechanism to reduce the number of effectors produced during the phase of clonal expansion.

**Figure 7:**
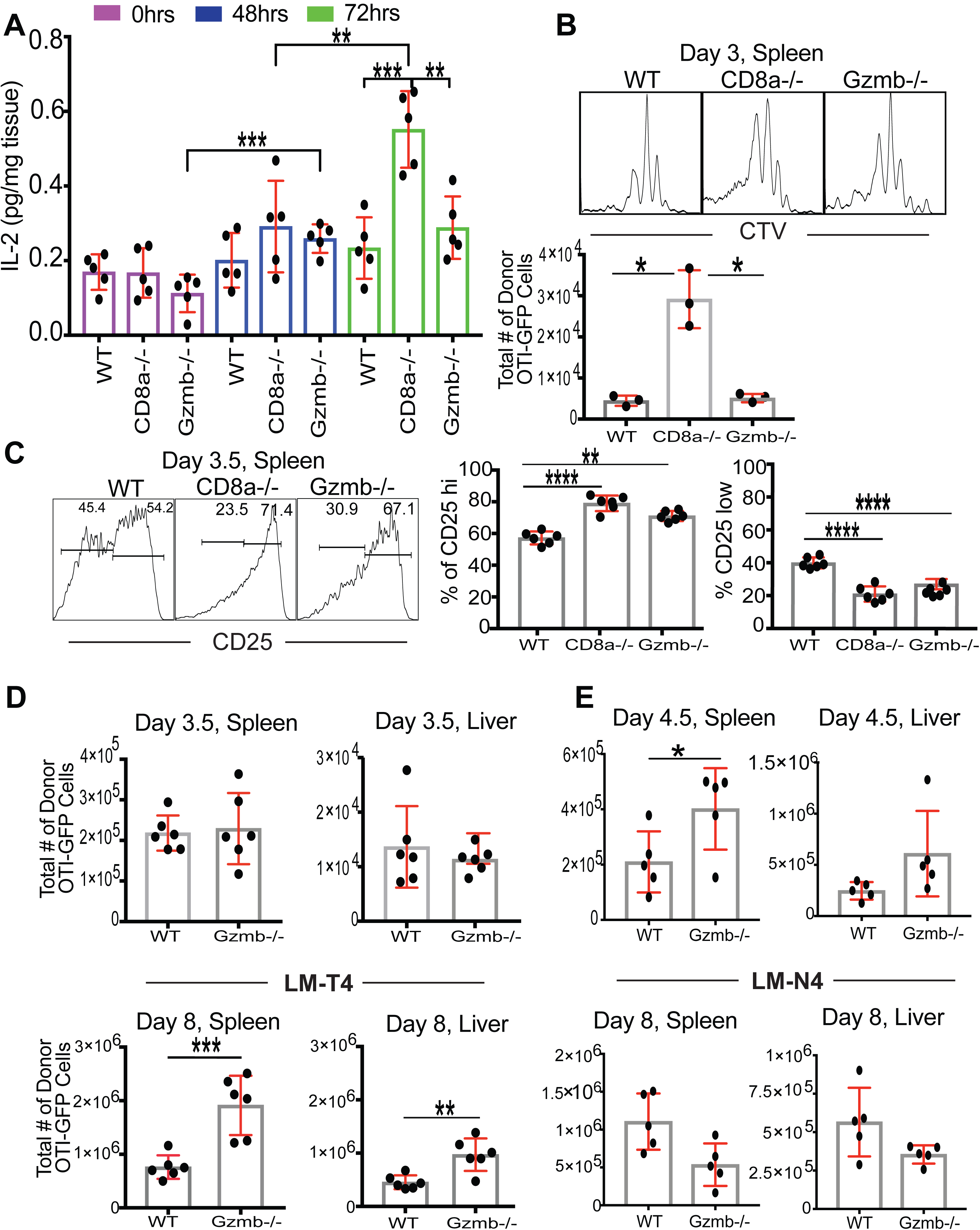
Reduced rate of clonal expansion of responding CD8 T cells due to limited duration of antigen presentation. A) CD8α -/- and *Gzmb -/-* mice have increased interstitial IL2 after infection with *Lm-ova*. CD8α -/- mice do not experience any decrease in the number of DCs due to the absence of CD8 T cells. *Gzmb -/-* mice experience minimal decrease in the number of DCs due to absence of PMC activity. Proliferation and differentiation of a small number of reporter OTI-GFP cells that were transferred prior to infection in B6, CD8α -/- and *Gzmb -/-* mice are shown in rest of the panels. *Lm-ova* expressing the T4 APL (lowest affinity) is used in panels B to D whereas *Lm-ova* expressing the natural N4 APL (highest affinity) is used in panel E. B) Increased rate of proliferation of reporter OTI-GFP cells in CD8α -/- mice. Due to the absence of pre-mitotic cytotoxicity, OTIs undergo additional rounds of divisions in CD8α -/- mice, as evidenced by the CTV dilution histograms. Accordingly, ~7-fold more reporter cells are present in the spleen of CD8α -/- within the first 3 days after infection. C) More of the responding OTIs show very high expression of CD25, the α subunit of IL2 receptor, in CD8α -/- and *Gzmb -/-* mice 3.5 days after infection. Representative histograms and quantifications for all mice are shown. The extent of increase in the % of CD25hi cells mirrors the extent of increase in the duration of antigen presentation in *Gzmb -/-* and CD8α -/- mice. D) Absolute counts of reporter OTI-GFP cells in B6 and *Gzmb -/-* mice 3.5 (top) and 8 (bottom) days after infection. The absolute counts of OTIs at 3 days (in B) and 3.5 days post-infection reveals that OTIs undergo much less net expansion after the first 3 days in CD8α -/- mice and then crash completely by 8^th^ day. ~2 fold more cells after 8 days would mean an additional round of division in *Gzmb -/-* mice compared to wild-type B6 mice. E) Absolute counts of reporter OTI-GFP cells in B6 and *Gzmb -/-* mice 4.5 (top) and 8 (bottom) days after infection with *Lm-ova* expressing the N4 APL. Each data point in the figure represents a separate mouse. The bars represent mean values. Standard deviation is shown in red. Standard deviation is shown in red. Statistical significance of difference in values was calculated by paired t test. p values from two-tailed tests are denoted as follows in the figures: *p < 0.05, **p < 0.01, ***p < 0.001, ****p < 0.0001.

## Discussion

Accumulating evidence on pre-mitotic cytotoxicity now allows us to consider it as a universal phenomenon during the priming phase of any CD8 T cell response. It has now been demonstrated in the context of *L. monocytogenes* (this study), LCMV (this study), Vaccinia Virus (Chiu et al., 2007) infection and implied in the context of malaria infection (Sano et al., 2001). Even very low affinity antigens can efficiently give rise to PMCs (Figure 1G and 1H), underscoring the breadth of the phenomenon. PMCs kill most of the conventional DCs (Figure 5A) which results in complete termination of antigen presentation within the first three days of *L. monocytogenes* infection (Figure 6A). It must be borne in mind that signalling pathways promoting necroptosis and pyroptosis are also operating in mature DCs during infection (Man and Kanneganti, 2016; Vandenabeele et al., 2010). Factors expressed by intracellular pathogens and type I interferons can also cause cell death (Carrero et al., 2004). Yet, in granzyme-B deficient mice most of the DCs remain viable in the first 48 hours of infection (Figure 5D). Thus, pre-mitotic cytotoxicity is the predominant mechanism by which DCs die early on during infection, although perhaps potentiated by other means.

*L. monocytogenes* infection in the spleen gives rise to ~5×10^4^ PMCs (Figure 4D) and >6×10^5^ conventional DCs (Figure 5A) die off within the first 48 hours. Numerically, this is a remarkable feat of serial killing by PMCs, as they have to act in an antigen-specific manner (Figure 1 and 5F). Pre-mitotic cytotoxicity develops during the later stage of priming, when the responding CD8 T cells are in contact with DCs typically for 10-30 minutes (Mempel et al., 2004). Thus, it is more likely that PMCs kill DCs by cooperative killing (Halle et al., 2016). Mature DCs secrete multiple chemokines to attract T cells to promote recognition by antigen-specific T cells, which should also facilitate cooperative killing (Castellino et al., 2006). Expression of granzyme B is very high in PMCs (Figure 1B, Figure 2B). This presumably allows them to keep releasing some fraction of granzyme B into engaged DCs and then quickly move to some other DC presenting antigen. Thus, DCs themselves likely receive granzyme B from multiple PMCs leading to cooperative killing.

One the other hand, finding 5 x 10^4^ PMCs for a single pathogen may be surprising in light of the observation that there are only 20-1000 antigen-specific naïve T cells per mouse for immune-dominant antigens (Jenkins and Moon, 2012). However, TCR recognition has been characterized as degenerate, with a single TCR recognizing hundreds of peptides with high affinity (Birnbaum et al., 2014). Our observation that lower potency pMHC, that induce poor clonal expansion (Zehn et al., 2009), were effective at generating PMCs (Figure 1H) together with this degeneracy may explain the breadth of the PMC response. This compares in scale to some aspects of innate immunity. We speculate that many PMC will not contribute to the clonally expanded population of immunodominant T cells, but nonetheless participate in regulation of the response and early pathogen control in the DC niche.

Actively dividing CD8 T cells do not make contacts with antigen presenting DCs (Bohineust et al., 2018) and once the division starts, they divide every 4-6 hours (Zhang and Bevan, 2011). Thus, PMCs are able to release their granzyme B into DCs only during the pre-mitotic phase. A very brief period of pre-mitotic cytotoxicity likely results in transient impact on the DC population. Coupled with the fact that lymphoid resident DC subsets have a rapid turnover rate (half-life of ~3 days) (Kamath et al., 2000), we expect antigen presentation by DCs to gradually resume as long as the pathogen persists beyond the first 3 days or so. The time-lag between pre-mitotic cytotoxicity and elimination of most of the DCs is presumably accounted for by the need for sufficient granzyme B to enter into the cytosol from vesicular compartments that can overcome the inhibition by Spi6 and then carry out proteolysis of a critical mass of substrates to irreversibly trigger apoptosis (Albeck et al., 2008). How granzyme B enters the cytosol from endosomes or analogous compartments in a perforin-independent manner is not known. We speculate that mechanisms that mediate cross-presentation facilitate the entry of internalized granzyme B into the cytosol of DCs.

Despite their apparent serial killing capacity in the context of lymphoid resident DCs in situ (Figure 5A), PMCs appear to be effective against a limited set of target cells (Figure 3A) and are considerably less potent compared to the conventional effector CTLs (Figure 3A-C). Increased expression of perforin (Figure 1A), expression of additional cytotoxic molecules such as fasL, granzyme A and possibly other mechanisms contribute to the high-potency of conventional CTLs (Janssen et al., 2010). Conventional effector CTLs accumulate cytolytic granules during differentiation through a gradual process of granule biogenesis and maturation over many days (Olsen et al., 1990). It is possible that our procedure for staining does not access all of granular granzyme B in the effector CTLs owing to dense packing of proteins in the granules. This possible caveat could have led to some degree of overestimation of the killing potency of effector CTLs in comparison to that of PMCs (Figure 3A-C).

Several pathogens use DCs to establish the infection and/or disseminate further into the host (Table S1). As PMCs act quite early during infection, granzyme B released into the cytosol of DCs can have a direct bactericidal effect to limit further expansion (Dotiwala et al., 2017). Elimination of DCs by PMCs can further serve to control the infectious burden in the lymphoid organ (Figure 6D) and limit dissemination. Elimination of DCs by PMCs may reduce immunopathology, but it leaves the host vulnerable to a secondary infection, as resident DC’s presentation of antigen to T cells is nearly absent immediately after the first infection (Figure 6C). Recently, it was shown that the continuous layer of macrophage lining the sub-capsular sinus in lymph nodes is disrupted by the DCs migrating from the periphery. This disruption hampers presentation of antigens to B cells and thus undermines B cell response to a secondary infection (Gaya et al., 2015). In addition, a primary infection undermines several aspects of innate immunity allowing an easy route for opportunistic pathogens (Hendaus et al., 2015). Thus, multiple mechanisms, including pre-mitotic cytotoxicity, contribute to a very high prevalence of secondary infections (Table S2).

Elimination of DCs by PMCs reduces and then terminates antigen presentation. Transient nature of antigen presentation, despite the prolonged presence of the pathogen has been documented before (Hafalla et al., 2003; Wong and Pamer, 2003). In fact, these studies also implicated the already activated CD8 T cells in a negative feedback role to limit antigen presentation. In this study, we have demonstrated that PMCs kill most of the conventional DC subsets in a granzyme B-dependent and a largely perforin-independent manner within two days (Figure 3). Termination of antigen presentation within three days (Figure 6A) in turn limits the clonal expansion of antigen-specific CD8 T cells already recruited to the response (Figure 7). It also prevents recruitment of additional antigen-specific naïve CD8 T cells that enter the lymphoid organ later during the response (Figure 6A). We expect these findings to extend to the CD4 T cells as well. In our opinion, limiting the pool of effector cells generated during the phase of clonal expansion by auto-inhibitory feedback, is the primary function of pre-mitotic cytotoxicity. Effector cells provide sterilizing immunity to infection but also cause tissue damage in the process and carry the potential to set off autoimmune response (King et al., 2012). Thus, restraining the number of effector cells produced is beneficial to the long-term survival of the host. We note that effector CTLs also perform the function of killing rare antigen-presenting cells later during the immune response. In fact, knock-out models with perforin deficiency in T cells and Fas deficiency in DCs develop chronic inflammation and autoimmunity due to persistent antigen presentation and T cell activation (Chen et al., 2012). This also mirrors the syndrome of hemophagocytic lymphohistiocytosis (HLH) in people with mutations that lead to defects in perforin-dependent cytotoxicity (Terrell and Jordan, 2013). Thus, limiting antigen presentation to a short duration is critical for the host. Pre-mitotic cytotoxicity uniquely contributes via a largely perforin-independent mechanism to nearly terminate antigen presentation within three days of an acute infection. Overall, our study demonstrates that adaptive immunity can exert its influence without any clonal expansion.

### Experimental Procedures

All experimental procedures are detailed in the supplementary material. The supplementary file also includes three additional figures and two tables.

### Author Contributions

DBD conducted the experiments with intermittent support from VM, DAB and RDDNA. VM conceptualized the project. VM and DBD designed experiments and analysed results. MLD, VM, and JJL shared research supervision. VM, MLD, and DBD interpreted the results and prepared the manuscript. All authors read and approved the manuscript.

## Acknowledgement

Supported by US National Institutes of Health grant R37 AI43542 (to MLD) and Wellcome Trust Principal Research Fellowship 100262/Z/12/Z (to MLD). Assistance of the Flow Cytometry and Light Microscopy Core Facilities at the NYU Medical Center is also acknowledged. We thank many colleagues for mice and reagents used in the study. We thank G. Lauvau, K. Khanna and A. Gerard for comments on the work.

